# Bell Jar: A Semi-Automated Registration and Cell Counting Tool for Mouse Neurohistology Analysis

**DOI:** 10.1101/2022.11.09.515722

**Authors:** Alec L.R. Soronow, Matthew W. Jacobs, Richard G. Dickson, Euiseok J. Kim

## Abstract

To investigate the anatomical organization of neural circuits across the whole brain, it is essential to register the experimental brain tissues to a reference atlas accurately. This procedure is also a prerequisite to quantify the locations and numbers of cells of interest in specific regions. However, it remains challenging to do registration on experimental tissue due to the intrinsic variation among the specimens, tissue deformation introduced by histological processing, and the potential inconsistency in the judgment of the experimenter during manual annotation. Here, we introduce Bell Jar, a multi-platform analysis tool with semi-automated affine warping of atlas maps onto microscopic images of brain slices and machine learning-based cell detection. Bell Jar’s intuitive GUI and internal dependency management enable users of all skill levels to obtain accurate results without programming expertise. To compare the performance of Bell Jar with previously published methods^**1–4**^, we labeled neurons in the mouse visual cortex with either an engineered rabies virus or an adeno-associated virus (AAV) for neural circuit tracing^**5**^, and quantified Bell Jar’s performance at each step of the pipeline for image alignment, segmentation, and cell counting. We demonstrated that Bell Jar’s output is as reliable as manual counting by an expert; it is more accurate than currently available techniques even with noisy data and takes less time with fewer user interventions. Bell Jar is an easy-to-navigate application that provides a reproducible, automated analysis workflow to facilitate the precise mapping of histological images of the mouse brain to the reference atlas and the quantification of cellular signals it is trained to recognize.

## Introduction

The quantitative analysis of brain tissue histology plays an important role in understanding the spatial organization of various neural cell types and their input-output connectivity across the brain. A typical histology pipeline includes sectioning the brain at a specific angle (*e*.*g*., coronal or sagittal), staining (*e*.*g*., immunofluorescence), mounting tissue sections on the slide, and microscope imaging. Inevitably, each brain differs somewhat in size and shape, and may be slightly deformed by tissue processing. Moreover, the sectioning angle may vary due to imperfect mounting of the brain on the sectioning apparatus (*e*.*g*., vibratome or cryostat). Consequently, precisely assigning labeled cells in brain slices to specific anatomical areas becomes challenging. Often, the experimenter manually draws area boundaries based on a reference atlas such as the Allen Mouse Brain Atlas^6^ or Paxinos and Franklin’s *The Mouse Brain in Stereotaxic Coordinates*^7^. However, this approach is highly subjective with personal biases and variations in judgments, which is further confounded by the discrepancy between the actual sectioning angle and that used for the atlas.

To improve the accuracy and consistency of histological image analysis, researchers have developed software to automate parts or all of the quantification procedure^1–4^, with promising results. However, some caveats have limited their use by the research community. The first barrier to adoption is the prerequisite computer expertise. Existing tools often consist of a collection of scripts^3,4^ rather than a centralized application, which makes them challenging to manage for people who are not computer-savvy. Users who are accustomed to intuitive graphic user interfaces (GUIs) may also find the command line tools cumbersome to use. The second barrier is that these tools are either fully automated or completely manual. A fully automated tool usually requires preprocessing and fine-tuning for each experiment, which poses challenges to labs working with a variety of tissue preparations. Moreover, as the automated pipeline can seldom guarantee absolute correctness given the complexity of experiments, proofreading and manual correction is almost always mandatory.

To address these limitations, we developed Bell Jar, a single cross-platform application that combines robust machine learning-based automatic processing and manual fine-tuning to provide a user-friendly and accurate quantification tool for neurohistology. The application helps users align and analyze images of brain slices in a directed workflow that minimizes user input. First, images of brain slices (“experimental images”) are automatically preprocessed to identify those without background objects or debris, and the user confirms the selection. Next, a convolutional autoencoder creates embeddings of the selected images. They are then compared with precomputed embeddings of the Allen Mouse Brain Atlas^6^ images over a range of sectioning angles (“reference images”) to find the angle that best matches the experimental images to the atlas. The user can then fine-tune the reference image predictions to optimize the alignment. Finally, the atlas images are warped to map them onto the corresponding experimental images, and a cell detection tool based on an object detection network called YoloV5^8^ is applied. This pipeline yields an accurate and reproducible assignment of anatomical areas to images of brain slices.

## Results

### Bell Jar’s architecture for cross-platform functionality

One of the primary goals of Bell Jar is to function across platforms. To accomplish this, we built the program’s GUI using Electron^9^ (Fig. 1). The Electron framework is based on open-source browser-based technologies so it can run on any machine that supports modern web browsers. Electron also allows bundling to native application packages on Windows, OSX, and Linux, so users can download the tool instead of building it from the source code. The compartmentalized architecture ensures correct dependencies regardless of the operating system (Fig. 1). While Python scripts execute the app’s functionality, the Electron web-app front-end provides a clean, easy-to-understand GUI for users to interact.

**Figure 1.**
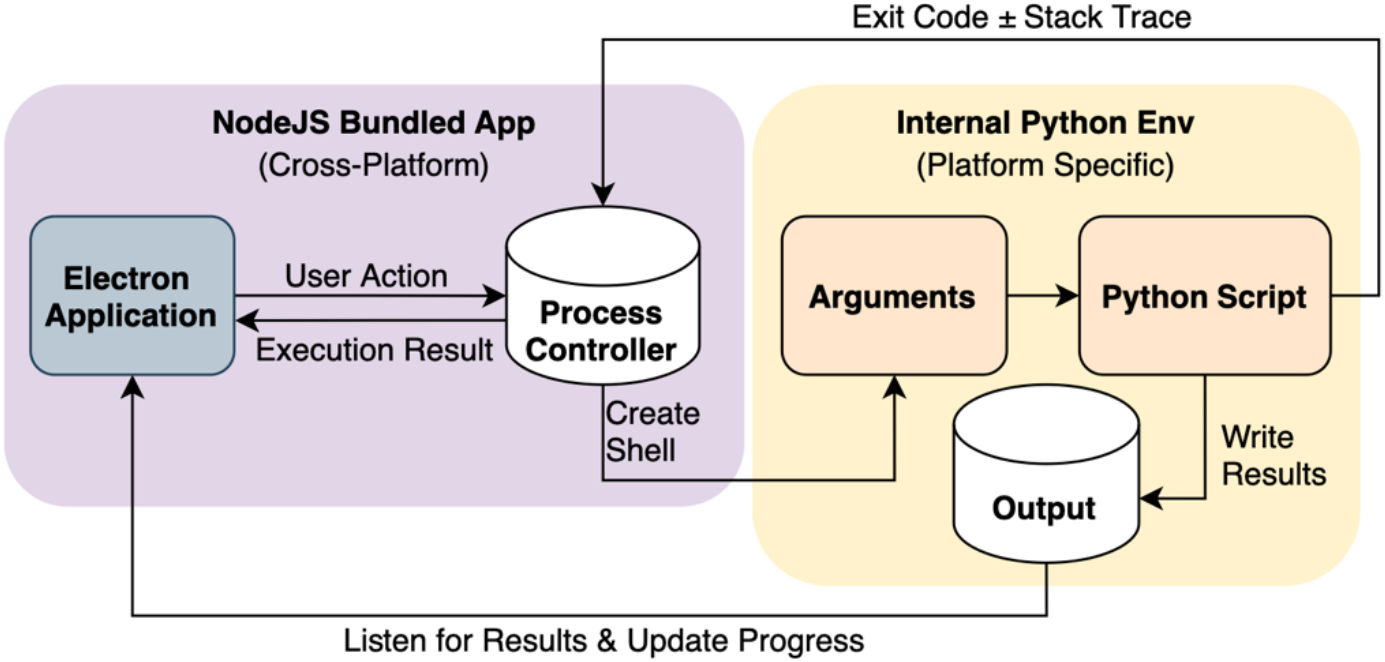
Bell Jar application architecture. The purple box contains the processes within the Electron framework, and the yellow box details the pathway within the Python environment. When a user selects an action in the program (i.e., detect cells), the relevant directory paths and arguments are collected and passed to a generated shell which executes the Python script for the selected task. The Python script also outputs status information, which the main Electron process listens to, updating the user about the task’s progress.

### Histological image alignment and registration

Bell Jar uses a convolutional autoencoder network (Fig. 2) to determine the tissue sectioning angle and the correspondence between experimental images and the atlas. As we intend to have Bell Jar produce good results while still be able to run locally on a consumer computer equipped with GPU (see Hardware Specifications in Materials and Methods), we minimized the depth of the convolutions and used only one fully connected layer. We used PyTorch^10^ to build and train both the encoder and decoder networks. To train the encoder, two synthetic datasets of 25,584 atlas images of sections of whole brains or single hemispheres were generated by rotating the Allen Mouse Brain Atlas to simulate cutting at different angles^6,11^. The angles ranged from −10 to +10 degrees, which covers the most common cutting angles on a microtome or a cryostat. Once the encoders were trained sufficiently, they were used to create reference embeddings of all the atlas images from the synthetic datasets for future fast comparison with experimental images.

**Figure 2.**
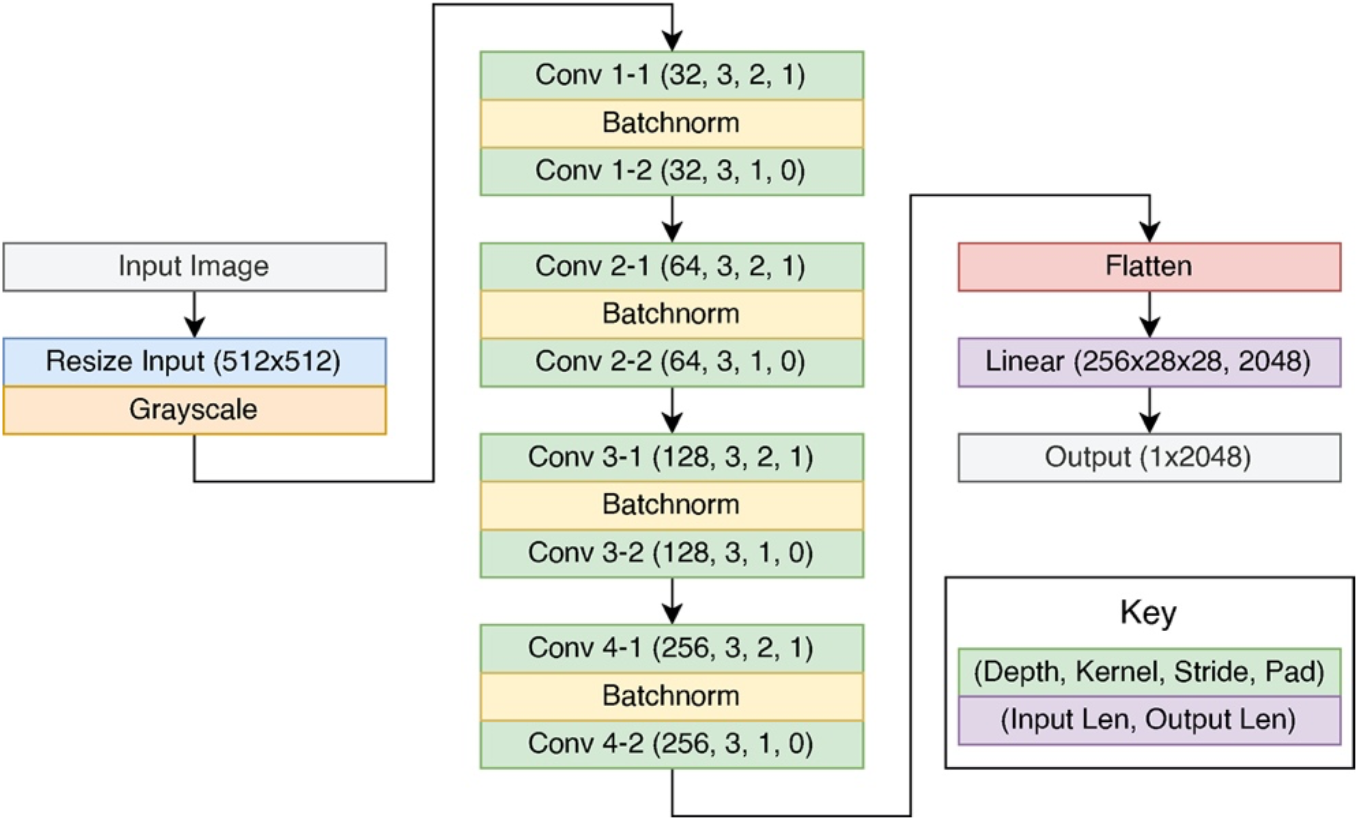
Architecture of the convolutional autoencoder for Bell Jar. Each input image is resized to 512 × 512 pixels and fed into the autoencoder to generate a unique embedding. The image passes through a series of increasingly deep convolutions; the output is flattened into a vector, which then runs through a fully connected layer to produce the output embedding. The output embedding uniquely identifies the image and its features, facilitating subsequent comparisons with the atlas.

During alignment the encoder network transforms each experimental image into its vector embedding, which is then compared with the embeddings of rotated reference images to determine the best match by minimizing the cosine distance (Fig. 3). After all experimental images are matched to the reference atlas, the majority sectioning angle (the angle occurring most frequently in the top matches for experimental images) is taken as the true cutting angle (Fig. 3A). An override is included for cases in which poor tissue quality hampers cutting angle prediction, or the cutting angle is already known precisely. Once the true angle is found, a final search step finds the best matching reference image (restricted to those sliced at the true angle) for each experimental image (Fig. 3B). The user can further fine-tune the matches to improve the alignment (Fig. 3C). The user navigates the experimental images and the reference image predictions from the autoencoder in one window. As the user adjusts the predictions to match the experimental images the average distance (number of sections apart) between each adjusted prediction is computed. We use the average distance to adjust all the predictions so that no prediction is further apart from another than the average distance.

**Figure 3.**
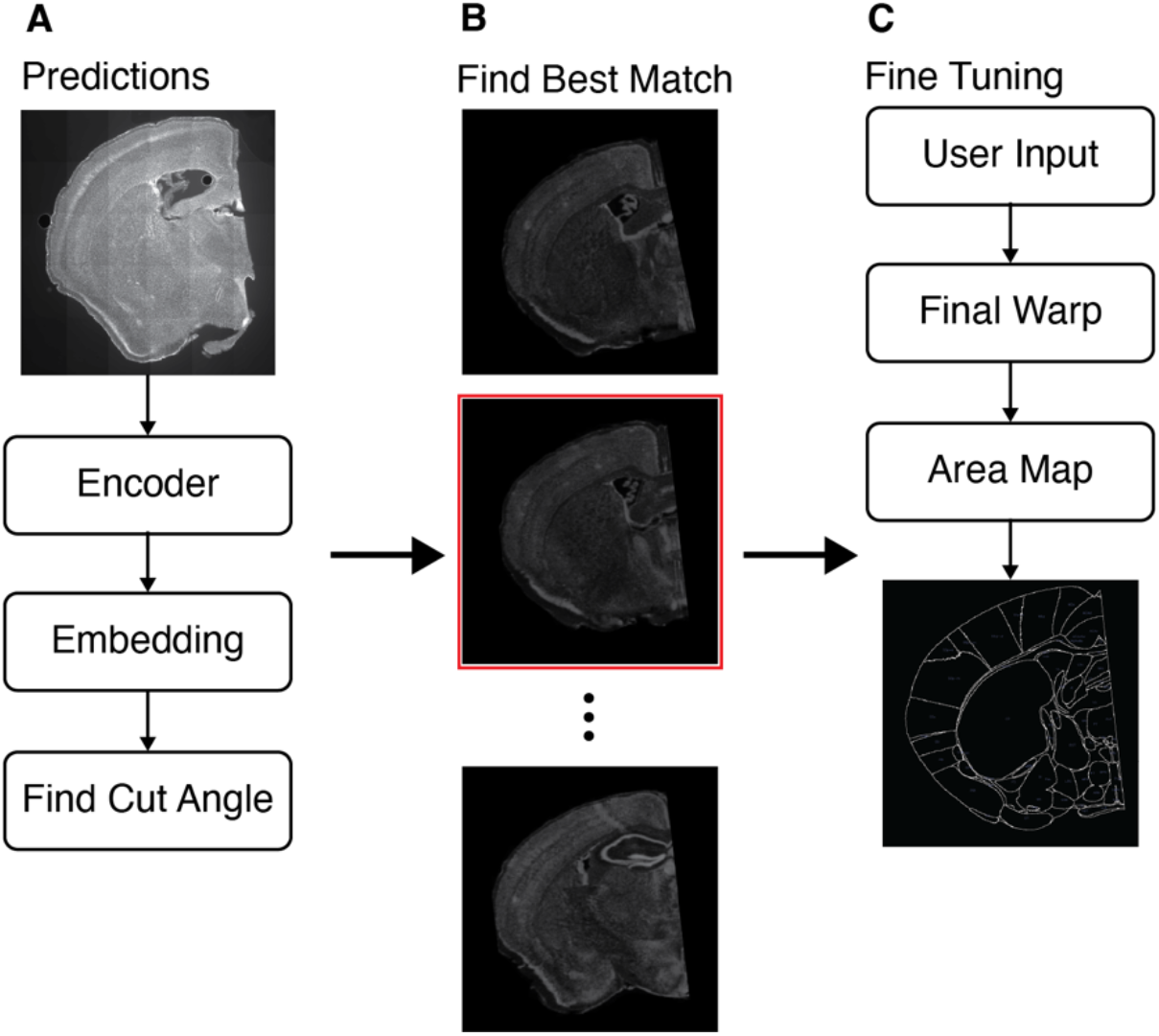
A single image example of the alignment and registration process. **(A)** The experimental image is passed to the encoder which transforms it to an embedding. The experimental image embedding is compared to the atlas embeddings to find its cut angle. **(B)** Restricted to within the cut angle a second pass of embedding comparison is run to find the best initial matching reference image. The best batch in this case is highlighted in red. **(C)** Users adjust the reference image predictions to best match the experimental image. When done the warping step occurs and the final registration is output.

Once all experimental images are matched to the atlas, Bell Jar performs registration using OpenCV and Numpy^12,13^. First, the reference image is resized to that of the experimental image. Then the contour of the brain slice is found in the experimental image and the reference image using a Gaussian filter and Otsu’s method^14^. The minimum rectangular bounding box for each contour is computed with OpenCV^12^. Finally, the reference image is warped onto the experimental image with a stepwise affine transformation. Specifically, the contents of the minimum bounding box in the reference image undergoes an affine transformation to fit it to the minimum bounding box of the experimental image. The resulting change is replicated on the reference images accompanying annotation files which label the regions of the reference section.

### Region-specific cell detection and quantification

To locate cells in the experimental images, we use YoloV5^8^, a lightweight object detection model adaptable to custom datasets. The detection step uses SAHI (Slicing Aided Hyper Inference)^15^ in combination with the YoloV5 model to detect the small neuronal cell bodies in the large images. Detected objects are saved to files along with the original image size to ensure proper scaling of cell coordinates during quantification of the cell counts per region in an experiment. The locations of the experimental cells must be recorded and integrated with the warp to quantify the number of cells per region in the experiment. To train our detector experimental images were collected from several experiments at random and screened for quality. Each image was flattened by maximum intensity projection, and tiles (640 × 640 pixels in size) where cells were present were collected. The resultant tiles were then labeled by hand to identify all neurons in each tile. The initial detector was trained on a total of 500 labeled neurons. After the rough detector had been trained, it was used to gather a superset of cells from all the experiments that the initial images were selected from. A custom script sliced each image into tiles and ran the rough detector to generate labels for neuron objects with 0.95 confidence and store each tile. After this process a new detector was trained with the newly created set of ∼8600 instances of neurons (Fig. 4). All trainings were done on the same GPU with a batch size of 16, using the Adam optimizer, and default YoloV5 hyperparameters^8^.

**Figure 4.**
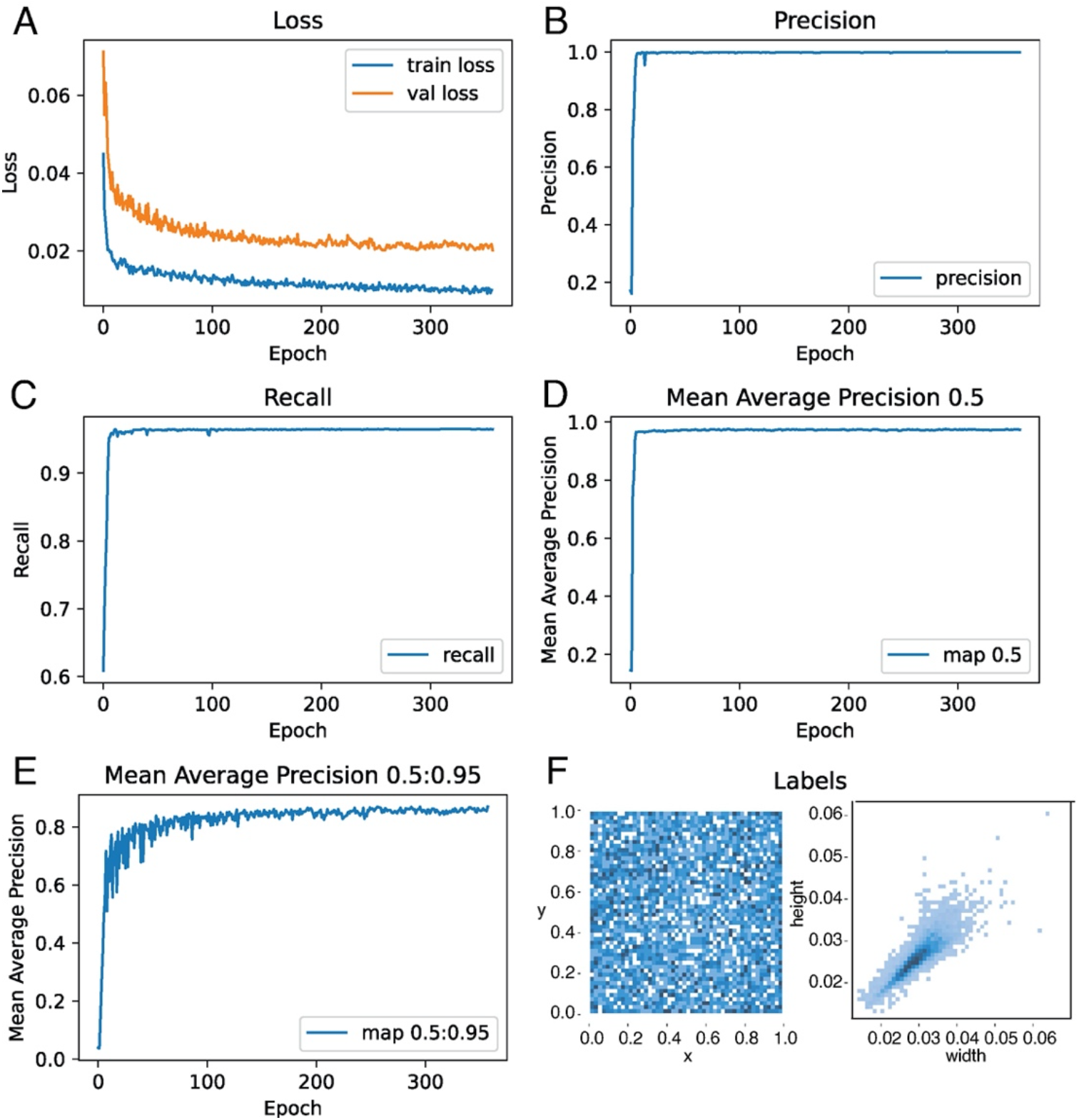
Performance metrics of the object detector training. **(A)** The validation and training loss of the detector. **(B)** The precision of the detector on the dataset. **(C)** The recall of the detector on the dataset over the epochs of training. **(D)** The mean Average Precision (mAP) of the detector at 0.5 confidence score. **(E)** The mAP of the detector for objects with 0.5 to 0.95 confidence. **(F)** Left: example matrix showing how often a labeled object was represented at a specific position in the dataset as normalized coordinates, showing the representative nature of the training set. Right: another matrix shows how often a specific size of object was represented in the training data in proportion to image size. Total training epochs = 357.

The quantification step takes data from the aforementioned steps and integrates them to provide experimental results. By reading the warped annotations, brain regions at any point in the experimental tissue can be mapped to a pixel coordinate. Using the cell locations found by the detector and iterating over the annotations provides a complete quantification. Once all sections are processed, the results per section and the whole experiment are written to a CSV file for study.

### Performance of the Bell Jar pipeline

With our new process the total time to completely quantify an experimental brain can be as little as one hour (Fig. 5). The primary time savings from the application has been the advances in cell detection using YoloV5 and SAHI^8,15^. Previous methods relied on image segmentation to process their cell images and obtain the locations of neurons^1,3^. Standard tools used in segmentation take significantly more processing power and memory than processing images in the Bell Jar pipeline. For example, to properly segment our images in the QUINT workflow^1^ using ilastik^16^, the images were down-sampled 30% or more to fit in memory during segmentation. Down sampling images lead to a loss in precision of cell location and clumping of neighboring cells. With the object detection framework in Bell Jar, these are not issues since inference is run on the original resolution images and does not rely on intensity thresholding to separate cells (Fig. 6).

**Figure 5.**
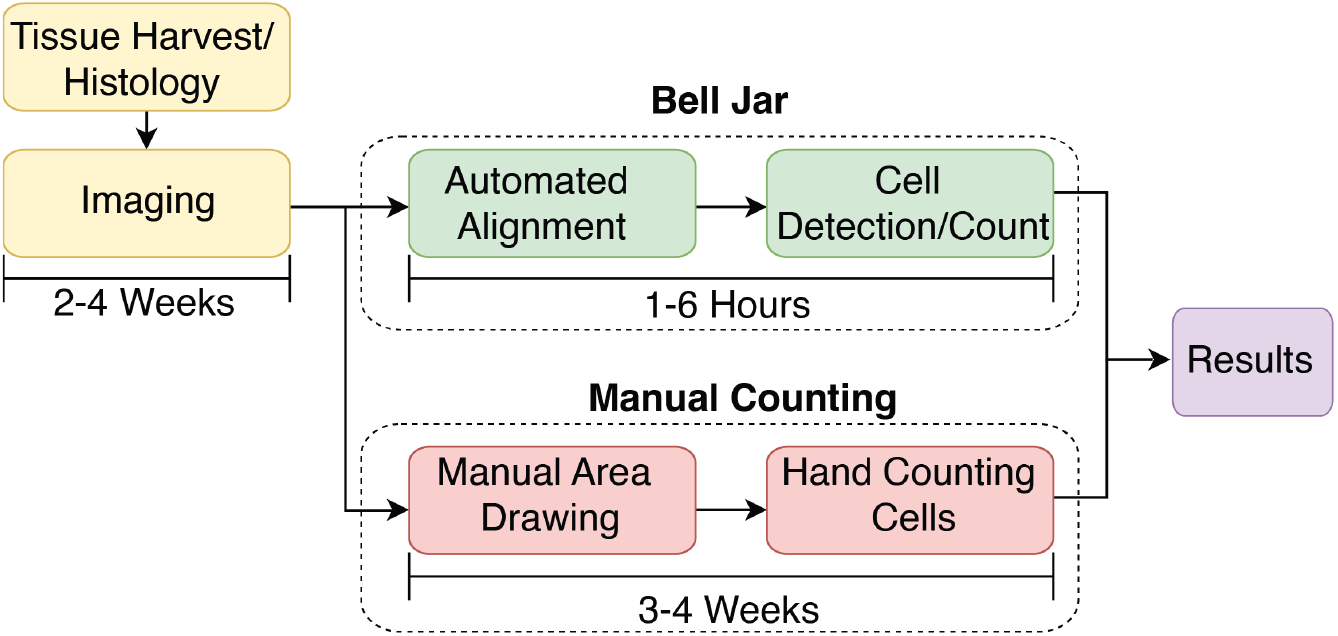
The Bell Jar workflow. The typical workflow of an experiment and the approximate time to results with Bell Jar versus manual counting. The initial surgery and imaging take weeks due to incubation times and scheduling. Therefore, it is crucial that the analysis be completed quickly to adjust downstream experiments and report on results. The DAPI images (nuclear stain) are used to align and warp the experiment to the Allen Brain reference atlas^6^. Afterward, fluorescent neuronal cell bodies are processed, and the cells ae detected with a YoloV5^8^ model using SAHI^15^. We transformed the coordinates of the detected cells into regions of the warped atlas for the finals results.

**Figure 6.**
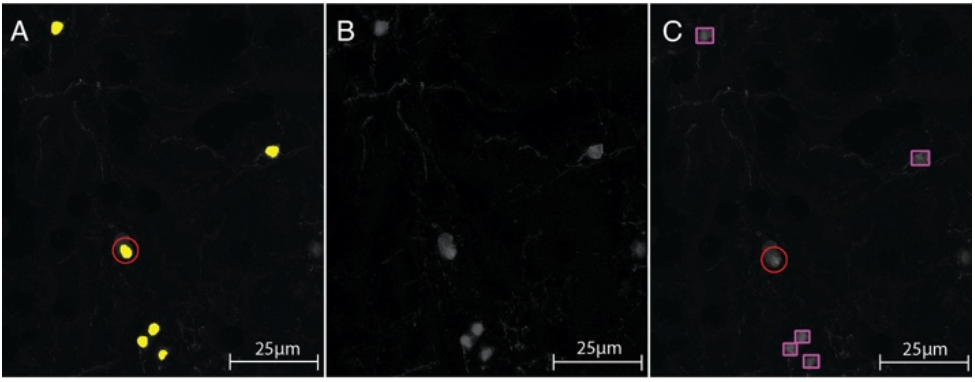
Comparisons of ilastik and Bell Jar for cell detection. **(A)** Yellow cells are segmented by ilastik^16^. Segmentation consists of a labeling step to select objects of the right intensity and a selection step to pick objects of the correct shape, which performed for each experiment in the ilastik software. Red circle: fluorescent debris incorrectly labeled as a cell. **(B)** The original pre-processed section of the tissue used in both these analyses, it has been filtered to even illumination and remove haloing from max intensity projection. **(C)** YoloV5^8^ trained on rabies virus-labeled neurons detects cells correctly (magenta rectangles) without erring on the debris (red circle).

To further test our method, we compared it against the segmentation and manual warping results that are typically used. We compared the alignments produced by Bell Jar’s pipeline against those of the QUINT workflow^1^ (Fig. 7). It was found that Bell Jar was better than QUINT^1^ in producing alignment maps that closely fit the original tissue section. The automated affine warping as opposed to manual adjustment enabled higher precision. Both the YoloV5^8^ detector and segmentation methods result in some loss of total counted cells as compared to manual counting (Fig. 8). However, each do so consistently across tissue samples. In this way, patterns can still be determined from the data collected from automated counting so long as precise raw counts are not of interest. Each method is strongly correlated to manual counting in terms of raw counts per section (Fig. 8). The two automated methods produce comparable results, the Bell Jar method benefits from a pre-trained object detector’s additional speed and ability to decern dimmer cells where thresholding may be incapable (Fig. 6). To validate that our object detection strategy could also be applied to other labeling types, we trained a second detector on nuclear specific labeling (scAAVretro-hSyn1-H2B-eGFP). In these tissue samples there were over a thousand cells per section on average. Nuclear specific labeling was chosen since high cell density, small signal area, and variation in intensity make counting these experiments with thresholding particularly challenging while simultaneously making manual counting tedious. Two sections were counted by hand for comparison to the new detector and thresholding based counts using ilastik^16^. Bell Jar’s object detection outperformed thresholding and was able to achieve performance similar to a human rater (Fig. 9). Object detection was able to pick up even dimmer cell objects that thresholding could not (Fig. 9D).

**Figure 7.**
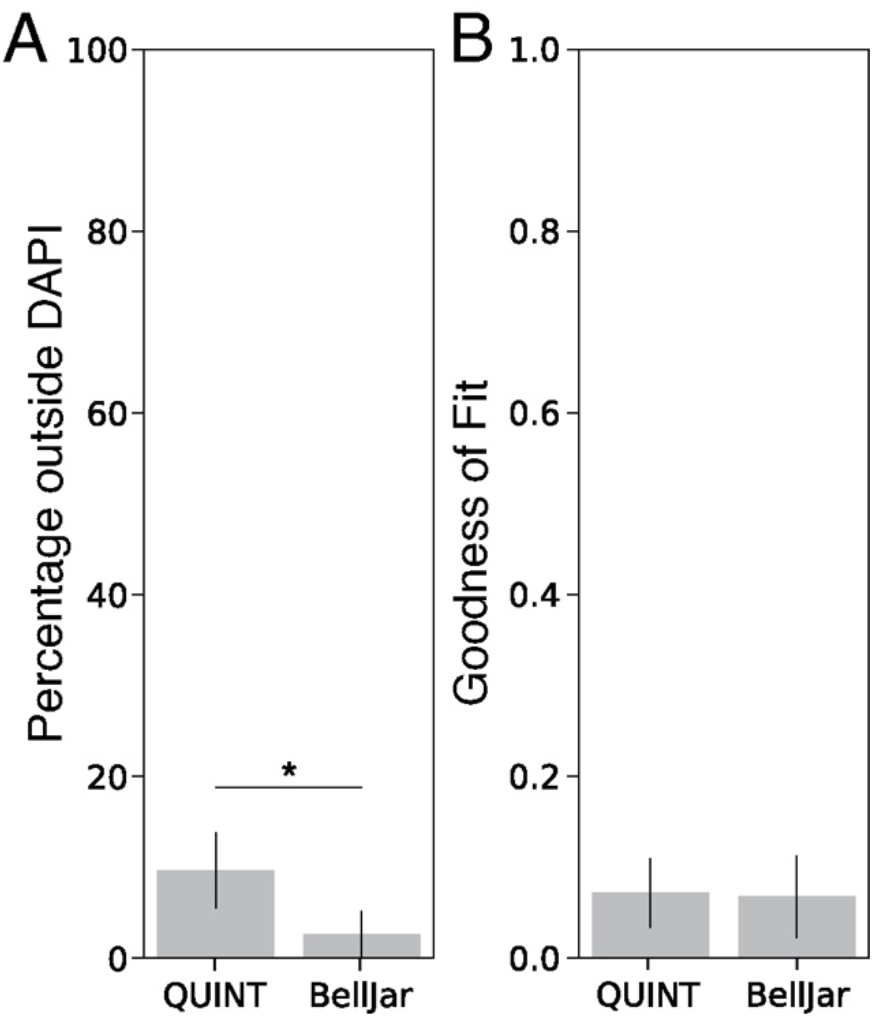
Measuring the quality of the semi-automated alignments. **(A)** A comparison between QUINT^1^ and Bell Jar alignments fit by comparing the overhang of their respective alignments. Bell Jar’s alignment uses a series of affine transformations to precisely warp the atlas information onto the tissue section. QUINT^1^ relies on the user to make these fine adjustments. Bell Jar alignment maps fit closer onto the original section (Student’s t-test, *, p<0.05). **(B)** A comparison between QUINT^1^ and Bell Jar alignment shape, lower values indicate closer fit. Each produces maps that closely resemble the given section.

**Figure 8.**
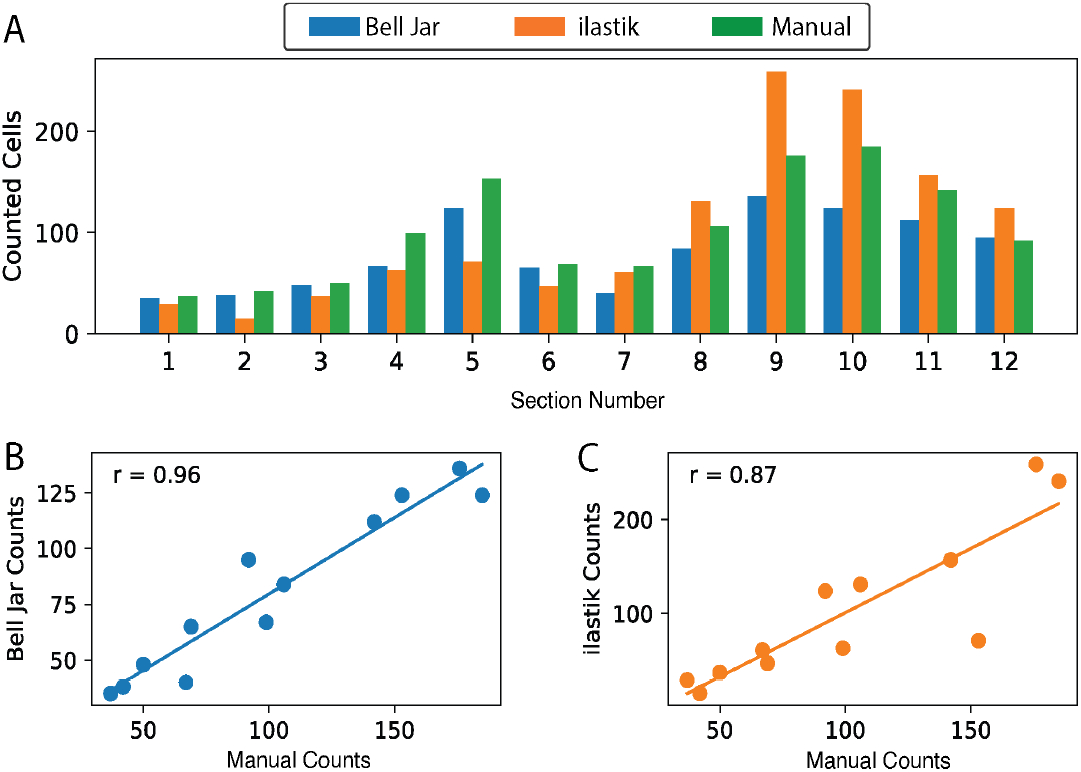
Comparing Bell Jar, manual, and thresholding-based counting. **(A)** Shown is a series of counts across twelve sections of tissue from a single animal. Each bar represents the total count of the method on the cells of that section. **(B)** A scatter plot showing the correlation of Bell Jar counts and manual counting. Bell Jar counts are strongly correlated (r=0.96) to manual counts. A scatter plot showing the correlation of ilastik^16^ counts and manual counting. Ilastik^16^ counts are also strongly correlated (r=0.87) to the manual observations.

**Figure 9.**
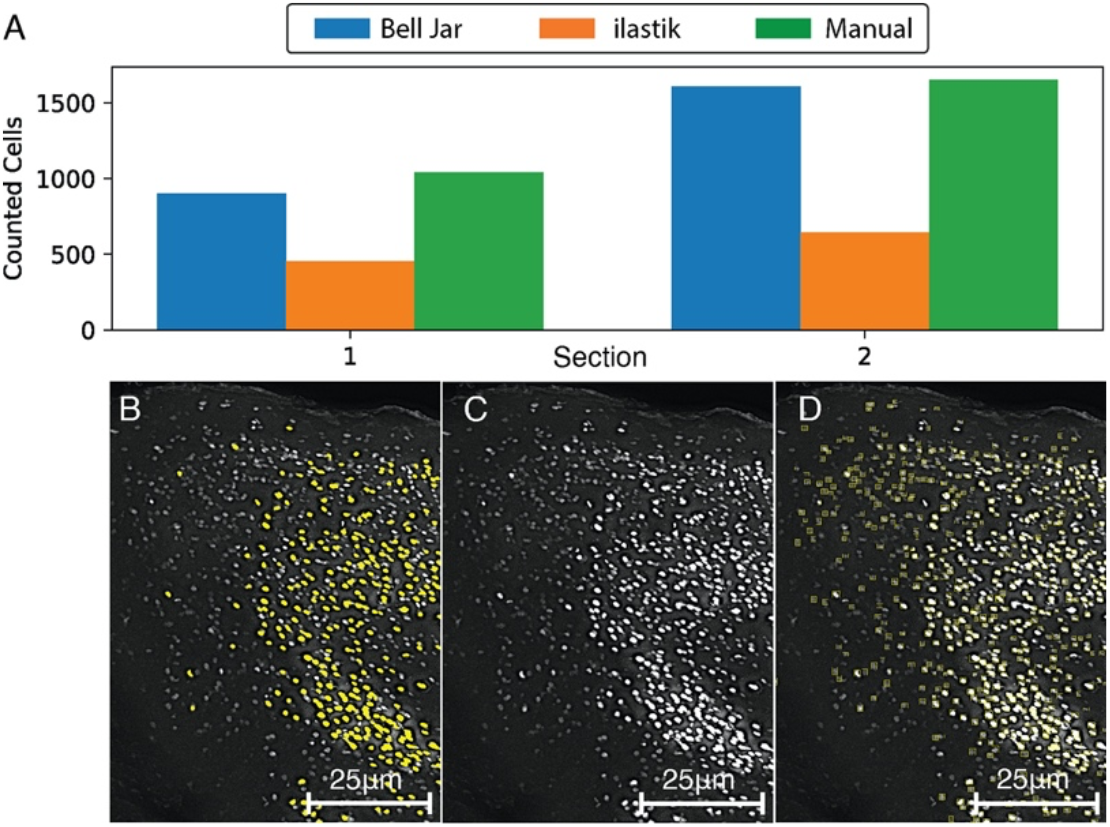
Comparing object detection and thresholding performance for high density counting. **(A)** A comparison of the counting results for two sections of nuclear labeled (scAAVretro-hSyn1-H2B-eGFP) cells among Bell Jar, ilastik^16^ and a human rater. **(B)** A zoomed in section of cells from section 2, in yellow are cells detected by thresholding with ilastik^16^. **(C)** The original zoomed in portion of section 2 with no overlay. A zoomed in section of cells from section 2, in yellow bounding boxes are cells detected by Bell Jar’s object detector.

## Discussion

Understanding the organizational principles of the mammalian brain at the cellular level is a fundamental but challenging task. It often requires tools to detect and quantify thousands of cells across different brain areas delineated according to standard atlases. Manual registration and quantification are time-consuming and laborious. Consistently integrating new automated techniques into the standard histology pipeline improves analyzing the volume of data produced by recent experiments. Using manual or previously published methods, aligning the microscope images of individual brain slices to a reference framework is challenging. Bell Jar provides the ability to register data to a standard reference atlas quickly and consistently enables faster turnarounds in the analysis of the underlying circuits. Its design enables a streamlined quantitative process that is easier for researchers to use and obtain conclusions from their data. Our automated affine transformation further reduces human error and provides a better-fitted registration. Additionally, Bell Jar allows for counting all regions of a hemisphere or whole brains where it is otherwise impractical by hand. Our tool reduces the burden of manual annotation allows experiments with high cell density or widespread tracing to be conducted and refined with high throughput. Using a lightweight model like YoloV5 also makes it easy to fine-tune the detector to work with other types of labeling, such as cytoplasmic or nuclear-specific fluorescent signals. Furthermore, results show that the method is superior to other automated counting methods and can serve as a tool to improve research workflows. The investigators can utilize Bell Jar to investigate a broad range of neuroscience questions, which include detecting c-fos staining to identify neurons activated during a particular behavioral task or detecting molecular markers for specific cell types in specific brain regions.

The Bell Jar suite relies upon cleaned input data like other methods and algorithms do. Microscopy data vary widely in brightness, distortions, debris, and tissue condition—these present challenges for one-size-fits-all tools. While the end user can clean most data for a workable analysis, some situations make analysis impractical such as tissue with extreme damage, stretching, or separated tissue (midbrain cleaved from posterior portion of cortex). Additionally, the method used for warping can be also subject to practical limitations. The estimated section, even with fine-tuning, cannot perfectly match the location of a given piece of experimental tissue in the brain. Performing an affine transformation on this non-exact match will further skew the resulting area definitions. Thus, it is not possible to reliably extract layer information (where misalignment skew results) from Bell Jar alignments. While these shortfalls are regrettable, they are present in all current automated solutions. Future research should address the need for sub-area level alignment, such as cortical laminar alignment.

A future study may extensively use Convolutional Neural Networks (CNNs) to predict layer information from patterns in DAPI staining accurately. A challenge in this regard is that the most reliable and widely used data set; the current version of Allen Brain Atlas has only Nissl volumes. Therefore, a custom dataset with accurately labeled DAPI-stained tissue would need to be created for training. Such a network could likely consistently outperform human raters. However, for now an adjustable alignment system with a human rater is the best option.

## Materials and Methods

### Experimental animals

GENSAT BAC transgenic SepW1-Cre NP39 mouse line has been previously described ^17^. C57BL/6J mice have been used as wild type. All experimental procedures followed protocols approved by the Institute Animal Care and Use Committee, University of California Santa Cruz.

### Virus preparation

All AAVs and EnvA+RVdG were produced by the Salk Viral Core GT3: scAAVretro-hSyn1-H2B-eGFP (AAVretro-nuclear eGFP, 1.32X10^13^ GC/ml), AAVretro-nef-lox66/71-tTA (1.77X10^12^ GC/ml), AAV8-TRE-DIO-oG-WPRE (5.92X10^12^ GC/ml), AAV8-TRE-DIO-GFP-T2A-TVA (7.00X10^13^ GC/ml), and EnvA+RV*dG*-mCherry (1.07X10^9^ Infectious Unit (IU)/ml).

### Animal surgery for virus injection

For rabies tracing experiments, SepW1-Cre NP39 mice received AAV helper injections at P87. Mice were anaesthetized with 100 mg/kg of ketamine and 10 mg/kg of xylazine cocktail via intra-peritoneal injections and mounted in a stereotax (David Kopf Instruments Model 940 series, Tujunga, CA) for surgery and stereotaxic injections. 50nl of AAVretro-nef-lox66/71-tTA was injected into the center of medial secondary visual cortex (V2M), using the following coordinates: 1.8 mm caudal, 1.6 mm lateral relative to lambda and 0.5-0.7 mm ventral from the pia. A 100 nl mixture of AAV8-TRE-DIO-oG-WPRE and AAV8-TRE-DIO-GFP-T2A-TVA was injected into the center of the primary visual cortex (V1), using the following coordinates: 3.4 mm caudal, 2.6 mm lateral relative to bregma and 0.5-0.7 mm ventral from the pia. We injected AAVs using air pressure by 1ml syringe with 18G tubing adaptor and tubing. To prevent virus backflow, the pipette was left in the brain for 5-10 minutes after completion of injection. Two or three weeks after AAV helper injection, 100 nl of EnvA+RV*dG*-mCherry were injected into the same site in V1 using 1ml syringe-mediated air pressure. Mice were housed for seven days to allow for trans-synaptic rabies spread and fluorescent protein expression.

For retrograde labeling experiments, C57BL/6J mice received 10 nl of scAAVretro-hSyn1-H2B-eGFP injection to V1 at P5. Mice were housed for three days to allow for retrograde AAV labeling and fluorescent protein expression.

### Histology and image analysis

Brains were harvested after transcardial perfusion using phosphate-buffered saline (PBS) followed by 4% paraformaldehyde (PFA). Brains were dissected out from skulls and post-fixed with 2% PFA and 15% sucrose in PBS at 4°C overnight, and then immersed in 30% sucrose in PBS at 4°C before sectioning. Using a freezing microtome, 50µm coronal brain sections were cut and stored in PBS with 0.01% sodium azide at 4°C. To enhance eGFP and dsRed signals, free-floating sections were incubated at 4°C for 16-48 hours with goat anti-GFP (1:1000; Rockland 600-101-215) and rabbit anti-dsRed (1:500; Clontech 632496) primary antibodies in PBS/0.5% normal donkey serum/0.1% Triton-X 100, followed by the appropriate secondary antibodies conjugated with Alexa 488 or 594 (Molecular Probes). Sections were counterstained with 10μM DAPI in PBS for 30 min to visualize cell nuclei. Immunostained tissue sections were mounted on slides with Polyvinyl alcohol mounting medium containing DABCO and allowed to air-dry overnight.

All sections were scanned with a 10x objective on a Zeiss AxioImager Z2 Widefield Microscope. Scanned images were first processed in Zeiss ZEN software using their extended focal imaging algorithm. Subsequent image files were processed and analyzed by Bell Jar, ilastik or NIH ImageJ (FIJI).

### Statistical Methods

For the analysis of the significance in the difference between the alignment methods fit and overlap Student’s t-test was used. For the calculation of the correlation of the various counting methods to manual data Pearson’s product moment correlation coefficient was used.

### Hardware Specifications

We did all training of any machine learning network described in the paper on a machine with a Threadripper 3960x processor and dual GTX 3090 GPUs operating in parallel. The minimum specifications we recommend for running Bell Jar are: an 8th generation Intel or Ryzen 5 (2017 onwards) processor and a GTX 1060 (6GB) GPU. The minimum GPU memory required for Bell Jar is 4GB, but we recommend at least 6GB for optimal performance. While a GPU is not strictly required to run Bell Jar, it is necessary to achieve the performance described in this paper.

## Data and Code Availability

We have already made the code for this project opensource, and it is freely available at https://github.com/asoronow/belljar for anyone to contribute. There are no releases in the limited repository now, and it must be built on your machine. An installer complete with auto-updates is planned.

## Acknowledgements

We thank A. Shariati and J. Lu for reading the manuscript, and P. Pham for mouse husbandry and histology. We acknowledge technical support from Benjamin Abrams, UCSC Life Sciences Microscopy Center, RRID: SCR_021135. We acknowledge support from the UCSC start-up fund, the Whitehall Foundation, the Hellman Fellows Program (E.J.K.), the Koret scholar program (A.L.R.S), and an NIH IRACDA Postdoctoral Training Grant (R.G.D.)

## References

1. Yates, S. C. et al. QUINT: Workflow for Quantification and Spatial Analysis of Features in Histological Images From Rodent Brain. Front. Neuroinformatics 13, 75 (2019).

2. Iqbal, A., Sheikh, A. & Karayannis, T. DeNeRD: high-throughput detection of neurons for brain-wide analysis with deep learning. Sci. Rep. 9, 13828 (2019).

3. Xiong, J., Ren, J., Luo, L. & Horowitz, M. Mapping Histological Slice Sequences to the Allen Mouse Brain Atlas Without 3D Reconstruction. Front. Neuroinformatics 12, 93 (2018).

4. Lauridsen, K. et al. A Semi-Automated Workflow for Brain Slice Histology Alignment, Registration, and Cell Quantification (SHARCQ). eneuro 9, ENEURO.0483-21.2022 (2022).

5. Kim, E. J., Jacobs, M. W., Ito-Cole, T. & Callaway, E. M. Improved Monosynaptic Neural Circuit Tracing Using Engineered Rabies Virus Glycoproteins. Cell Rep. 15, 692–699 (2016).

6. Wang, Q. et al. The Allen Mouse Brain Common Coordinate Framework: A 3D Reference Atlas. Cell 181, 936-953.e20 (2020).

7. Franklin, K. B. J. & Paxinos, G. The mouse brain in stereotaxic coordinates. (Elsevier Academic Press, 2008).

8. Jocher, G. et al. ultralytics/yolov5: v6.1 - TensorRT, TensorFlow Edge TPU and OpenVINO Export and Inference. (2022) doi:10.5281/zenodo.6222936.

9. Electron JS.

10. Paszke, A. et al. PyTorch: An Imperative Style, High-Performance Deep Learning Library. in Advances in Neural Information Processing Systems 32 (eds. Wallach, H. et al.) 8024–8035 (Curran Associates, Inc., 2019).

11. Virtanen, P. et al. SciPy 1.0: Fundamental Algorithms for Scientific Computing in Python. Nat. Methods 17, 261–272 (2020).

12. Bradski, G. The OpenCV Library. Dr Dobbs J. Softw. Tools (2000).

13. Harris, C. R. et al. Array programming with NumPy. Nature 585, 357–362 (2020).

14. Otsu, N. A Threshold Selection Method from Gray-Level Histograms. IEEE Trans. Syst. Man Cybern. 9, 62–66 (1979).

15. Akyon, F. C. et al. SAHI: A lightweight vision library for performing large scale object detection and instance segmentation. (2021) doi:10.5281/zenodo.5718950.

16. Berg, S. et al. ilastik: interactive machine learning for (bio)image analysis. Nat. Methods 16, 1226–1232 (2019).

17. Gerfen, C. R., Paletzki, R. & Heintz, N. GENSAT BAC Cre-Recombinase Driver Lines to Study the Functional Organization of Cerebral Cortical and Basal Ganglia Circuits. Neuron 80, 1368–1383 (2013).

